# Speed-dependent biomechanical changes vary across individual gait metrics post-stroke relative to neurotypical adults

**DOI:** 10.1101/2022.04.01.486769

**Authors:** Sarah A. Kettlety, James M. Finley, Darcy S. Reisman, Nicolas Schweighofer, Kristan A Leech

## Abstract

**Background:** Gait training at fast speeds is recommended to reduce walking activity limitations post-stroke. Fast walking may also reduce gait kinematic impairments post-stroke. However, the magnitude of speed-dependent kinematic impairment reduction in people post-stroke relative to neurotypical adult walking patterns is unknown.

**Objective:** To determine the effect of faster walking speeds on gait kinematics post-stroke relative to neurotypical adults walking at similar speeds.

**Methods:** We performed a secondary analysis with data from 28 people post-stroke and 50 neurotypical adults treadmill walking at multiple speeds. We evaluated the effects of speed and group on individual spatiotemporal and kinematic metrics and performed k-means clustering with all metrics at self-selected and fast speeds

**Results:** People post-stroke decreased step length asymmetry and trailing limb angle impairment, reducing between-group differences at fast speeds. Speed-dependent changes in peak swing knee flexion, hip hiking, and temporal asymmetries exaggerated between-group differences. Our clustering analyses revealed two clusters. One represented neurotypical gait behavior, composed of neurotypical and post-stroke participants. The other characterized stroke gait behavior, comprised entirely of participants post-stroke. Cluster composition was largely consistent at both speeds, and the distance between clusters increased at fast speeds

**Conclusions:** The biomechanical effect of fast walking post-stroke varied across individual gait metrics. For participants within the stroke gait behavior cluster, speed-dependent changes did not lead to an overall gait pattern more similar to neurotypical adults. This suggests that combining fast walking with an approach to strategically target gait metrics with smaller speed-dependent changes may potentiate the biomechanical benefits of fast walking.

## BACKGROUND

In the United States, approximately 795,000 people have a stroke each year [1], and gait dysfunction is a common outcome [2]. Two domains of gait dysfunction are kinematic impairments (e.g., increased circumduction and reduced knee flexion) [3,4] and activity limitations (e.g., decreased gait speed and independence) [5]. Kinematic and spatiotemporal impairments are associated with increased metabolic cost [6,7] and fall risk [8], whereas activity limitations, particularly gait speed, are associated with reduced community ambulation and quality of life [9]. Improvements in both categories are essential goals for people post-stroke and often targets during gait rehabilitation [10].

Recently, there has been an increased emphasis on structuring interventions to target activity limitations in walking post-stroke [11]. Current evidence suggests that the most effective way to do this is with moderate to high aerobic intensity gait training, often achieved by walking at faster speeds [5]. The shift away from interventions that prioritize reducing kinematic gait impairments post-stroke may be due, in part, to evidence that select kinematic metrics improve while walking faster. Specifically, people post-stroke reduce step length asymmetry and increase the magnitudes of paretic swing knee flexion and trailing limb angle with fast walking [12,13]. Furthermore, walking faster does not lead to increased circumduction – a common compensatory movement [12,14]. This suggests that improved post-stroke gait kinematics may be a byproduct of walking faster, even when not specifically targeted.

While walking faster can lead to changes in select kinematic metrics post-stroke relative to the habitual walking pattern, the magnitude of this speed-dependent change relative to neurotypical adults walking at similar speeds is unclear. That is, walking faster may reduce, exaggerate, or not affect the difference in post-stroke gait kinematics compared to neurotypical adults. Additionally, previous work has focused on speed-dependent changes in individual gait metrics post-stroke [12– 15], leaving the impact of fast walking on *overall* gait behavior post-stroke an open question.

This study evaluated the effect of gait speed on select spatiotemporal and kinematic gait metrics in people post-stroke relative to neurotypical adults walking at similar speeds. To do this, we performed a secondary analysis of three previously published data sets [7,12,16]. Based on studies that demonstrate abnormal muscle co-activation patterns post-stroke during walking [17– 19] and maximal hip extension torque production [20], we hypothesized that the difference between gait metrics between groups would increase as speed increases. We also evaluated all metrics simultaneously in a k-means clustering analysis to determine if the difference in overall gait behavior between people post-stroke and neurotypical adults changed with speed. We hypothesized that there would be two clusters (one that captured the gait behavior of neurotypical adults and one that captured post-stroke gait behavior) and that walking faster would cause the distance between the clusters to increase. This would indicate that the overall walking patterns of people post-stroke and neurotypical adults became more different at fast speeds.

## METHODS

### Participants

We performed a secondary analysis of data sets from three previously published cross-sectional studies of people post-stroke and neurotypical adults [7,12,16]. Specifically, data from people post-stroke were obtained from Tyrell et al., 2011 (n = 20) [12], data from people post-stroke (n = 15) and neurotypical adults (n = 15) from Finley and Bastian, 2017 [7], and additional data from neurotypical adults were obtained from Fukuchi et al., 2018 (n = 42) [16]. To evaluate continuous gait patterns, only data from participants whose slowest walking speed was >0.20 m/s were included. Participants without a standing calibration file were also excluded from these analyses. All data were de-identified; therefore, this analysis is not considered human subjects research and did not require review from the University of Southern California Institutional Review Board.

### Stroke Data

The post-stroke data set included lower extremity kinematic data of people ≥ six months post-stroke previously published in Tyrell et al., 2011 (n = 20) [12]. Participants walked on a treadmill at four different speeds in a randomized order: self-selected, fast-as-safely possible, and two intermediate speeds. The two intermediate speeds were chosen to be as equally distributed as possible between the self-selected and fast-as-safely possible speeds. Two participants only completed one intermediate speed; therefore, only three speeds were included in the analyses for these participants. Marker data were collected using a motion capture system sampling at 100 Hz. Details about the marker set can be found in the original publication [12]. Data were collected at each speed for two twenty-second trials, resulting in forty seconds of data.

The post-stroke data set also included data from participants published in Finley and Bastian, 2017 (n = 15) [7]. Participants walked at four speeds on a treadmill: self-selected speed, fastest possible speed they could maintain for five minutes, and 80% and 120% of their self-selected speed. One participant could not complete the trial at 120% of their self-selected speed; therefore, only three speeds were included in the analysis for this participant. The order of speed presentation was randomized, and participants walked on the treadmill for five minutes at each speed. For these analyses, the middle thirty seconds of each trial were analyzed. Marker data were collected using a motion capture system with infrared-emitting markers sampling at 100 Hz. Details about the marker set can be found in the original publication [7].

### Neurotypical Data

Part of the neurotypical data set included age- and speed-matched neurotypical adults to the people post-stroke in Finley and Bastian, 2017 (n = 15) [7]. Neurotypical adults walked on the treadmill at the same four speeds as the people post-stroke. The data collection procedures were the same as outlined above for the post-stroke group.

The age-matched neurotypical data set also included data from older adults from Fukuchi et al., 2018 (n = 18) [16]. Participants walked on a treadmill at eight speeds ranging from 40 – 145% (in increments of 15%) of their self-selected speed, in a randomized order. Our statistical analysis (described below) was not paired and did not require exact speed matching, so for each participant, we extracted four speeds that reflected a similar speed range to post-stroke speeds in the first stroke data set [12]. This ensured that all speeds in the stroke data set were represented in the neurotypical data set (Supplemental Table 1). Participants walked for ninety seconds at each speed, and data were recorded during the final thirty seconds. Marker data were collected using a motion capture system sampling at 150 Hz. Details about the marker set can be found in the original publication [16].

For the clustering analysis only, we included data from neurotypical young adults (n = 24) in addition to the post-stroke and neurotypical older adult data to make the analysis more robust. These data were collected in the same laboratory and manner as the older adults from Fukuchi et al., 2018 [16]. The data from young neurotypical participants were chosen to match the range of self-selected and fastest speeds seen in the first stroke data set [12].

### Data Analyses

We used spatiotemporal and kinematic metrics computed in Visual3D (C-Motion, Germantown, MD) from Tyrell et al., 2011 in all statistical analyses [12]. The marker data from the other data sets were processed and analyzed in MATLAB R2020a (MathWorks, Natick, MA). These data were low-pass filtered with a 6 Hz cutoff [21]. Foot-strike and toe-off were defined as the most anterior and posterior positions of the lateral malleoli markers, respectively.

The spatiotemporal and kinematic metrics of interest reported here were selected and defined to be consistent with those reported in Tyrell et al., 2011 [12] since we could not re-process those data. Spatiotemporal outcome measures were step length asymmetry, single-limb support time asymmetry, and double-limb support time asymmetry. Kinematic outcome measures were peak swing knee flexion angle, trailing limb angle, circumduction, and hip hiking. Markers used to calculate these metrics were iliac crest, greater trochanter, lateral femoral epicondyle, lateral malleolus, and fifth metatarsal. All intralimb kinematic outcome measures were calculated on the paretic (stroke) or the right limb (neurotypical). Mean values across all strides taken within the bin specified above were used in all statistical analyses.

### Statistical Analysis

To evaluate the effect of speed and group on the individual gait metrics, we fit robust linear mixed-effects models using the rlmer package [22] in R (4.0.2) [23]. We used a robust model instead of a traditional mixed-effects model to address normality and homoscedasticity assumptions violations. Fixed effects for group, speed, and speed by group interaction were included. Before fitting the models, we removed the mean from the speed values, which allowed us to interpret the group coefficient at the average speed across the sample. A random intercept term was included in all models to account for the repeated measures design. A separate model was fit for each outcome measure. P-values were calculated using the Satterthwaite approximation [24]. Statistical significance was set *a priori* at 0.05.

To capture the effect of speed on overall gait behavior, we used k-means clustering to identify subsets of participants with similar overall gait behavior and determined whether walking faster altered the composition of these subsets. We performed k-means clustering using the kmeans function in R (4.0.2) [23] with data from both groups at the self-selected and fastest speeds of the participants post-stroke using all seven gait metrics listed above. Because these metrics have different units, the data were scaled (mean = 0, standard deviation = 1) before clustering. We determined the number of clusters using the silhouette method. Within sum of squares was used to assess the between-subjects variability within each cluster and between sum of squares to determine the distance between clusters. To identify the relative importance of each variable in the determination of the clustering, we used the cluster assignment as the outcome variable in a random forest algorithm [25]. We then extracted the variable importance using the importance function in the randomForest package [25]. For each variable, the prediction error on the data that was not used to train the model was determined, then the prediction error when the data for each predictor variable was permuted (shuffled) was determined. The difference between the two prediction errors represents the mean decrease in accuracy for a specified variable; higher values represent greater variable importance [25,26]. We also performed a principal components analysis on the scaled data at both speeds to allow visualization of the clusters in two dimensions. Finally, we used Mann-Whitney U tests to determine if two key clinical measures of impairment, Lower-Extremity Fugl-Meyer scores and gait speeds, differed between the participants post-stroke who were assigned to different clusters.

## RESULTS

After combining data sets, 28 people post-stroke, 26 neurotypical older adults, and 24 neurotypical younger adults were included in these analyses. Eight participants were excluded due to their slowest walking speed being <0.20 m/s, and six were excluded due to a missing standing calibration file. For the robust mixed-effects analysis, we used post-stroke data (n = 28), neurotypical data from older adults obtained from Fukuchi et al., 2018 (n = 18) [16], and matched neurotypical controls from Finley and Bastian, 2017 (n = 8) [7]. For the clustering analysis, we included additional data from younger neurotypical adults (18 – 39 years; n = 24). Clinical demographics for participants post-stroke are included in Table 1.

**Table 1.** Clinical demographics of participants post-stroke.

| Data Set | Sex | Side of Paresis | Age (years) | Orthosis or Assistive Device | Lower-Extremity Fugl-Meyer | Speed 1 (m/s) | Speed 2 (m/s) | Speed 3 (m/s) | Speed 4 (m/s) |
| --- | --- | --- | --- | --- | --- | --- | --- | --- | --- |
| 1 | M | L | 66 | AFO | 18 | 0.80 | 0.90 | 1.10 | 1.30 |
| 1 | M | L | 50 | None | 19 | 0.90 | 1.00 | 1.10 | 1.20 |
| 1 | M | L | 74 | AFO | 17 | 0.30 | 0.40 | 0.50 | 0.70 |
| 1 | F | L | 59 | None | 31 | 0.70 | 0.80 | 0.90 | 1.00 |
| 1 | M | R | 61 | AFO | 14 | 1.00 | 1.10 | 1.20 | 1.30 |
| 1 | F | L | 66 | None | 21 | 0.40 | 0.50 | 0.50 | 0.60 |
| 1 | F | L | 78 | None | 24 | 0.70 | 0.80 | 0.90 | 1.00 |
| 1 | M | R | 57 | None | 15 | 0.60 | 0.80 | 0.90 | 1.10 |
| 1 | M | R | 75 | None | 31 | 0.70 | 0.90 | 1.10 | 1.30 |
| 1 | F | L | 51 | AFO | 20 | 0.40 | 0.50 | 0.60 | 0.70 |
| 1 | M | R | 47 | None | 25 | 0.80 | 0.90 | 1.00 | 1.10 |
| 1 | M | R | 60 | None | 25 | 0.90 | 1.00 | 1.20 | 1.30 |
| 1 | M | L | 52 | AFO | 20 | 0.80 | 1.00 | 1.20 | 1.40 |
| 1 | F | R | 62 | SPC | 25 | 0.50 | 0.60 | 0.80 | 1.00 |
| 1 | M | R | 72 | None | 32 | 0.80 | 0.90 | 1.00 | 1.20 |
| 1 | M | R | 71 | AFO, SPC | 19 | 0.70 | 0.80 | 0.90 | - |
| 1 | M | R | 77 | None | 22 | 0.70 | 0.80 | 0.90 | 1.10 |
| 1 | F | L | 45 | None | 23 | 0.90 | 1.00 | 1.10 | - |
| 1 | M | R | 73 | None | 31 | 0.70 | 0.80 | 1.00 | 1.30 |
| 1 | M | L | 78 | SPC | 23 | 0.50 | 0.60 | 0.70 | 0.90 |
| 2 | M | L | 42 | Not reported | 22 | 0.77 | 0.96 | 1.15 | 1.24 |
| 2 | F | R | 56 | Not reported | 33 | 0.46 | 0.57 | 0.68 | 0.81 |
| 2 | F | R | 68 | Not reported | 23 | 0.21 | 0.26 | 0.31 | 0.56 |
| 2 | M | R | 57 | Not reported | 17 | 0.48 | 0.61 | 0.72 | 1.00 |
| 2 | M | L | 52 | Not reported | 32 | 0.82 | 1.02 | 1.22 | 1.35 |
| 2 | M | L | 67 | Not reported | 27 | 0.26 | 0.33 | 0.40 | - |
| 2 | M | L | 54 | Not reported | 31 | 0.28 | 0.35 | 0.42 | 0.60 |
| 2 | M | L | 60 | Not reported | 33 | 0.22 | 0.28 | 0.34 | 0.57 |

Abbreviations: 1, Tyrell et al., 2011; 2, Finley and Bastian 2017; F, female; M, male; L, left; R, right; AFO, ankle-foot orthosis; SPC, single-point cane. Ankle-foot orthoses were worn during data collection. Single-point canes were not used during data collection but were used by participants during daily walking.

### Spatiotemporal Parameters

As expected [4], people post-stroke exhibited greater step length asymmetry (Figure 1A; β = 0.08, p < 0.001) compared to neurotypical adults. With increases in speed, neurotypical adults decreased step length asymmetry (β = -0.08, p < 0.001). However, people post-stroke decreased step length asymmetry more than neurotypical adults at faster speeds (speed by group interaction: β = -0.05, p < 0.001), reducing the difference between the groups at faster speeds.

**Figure 1.**
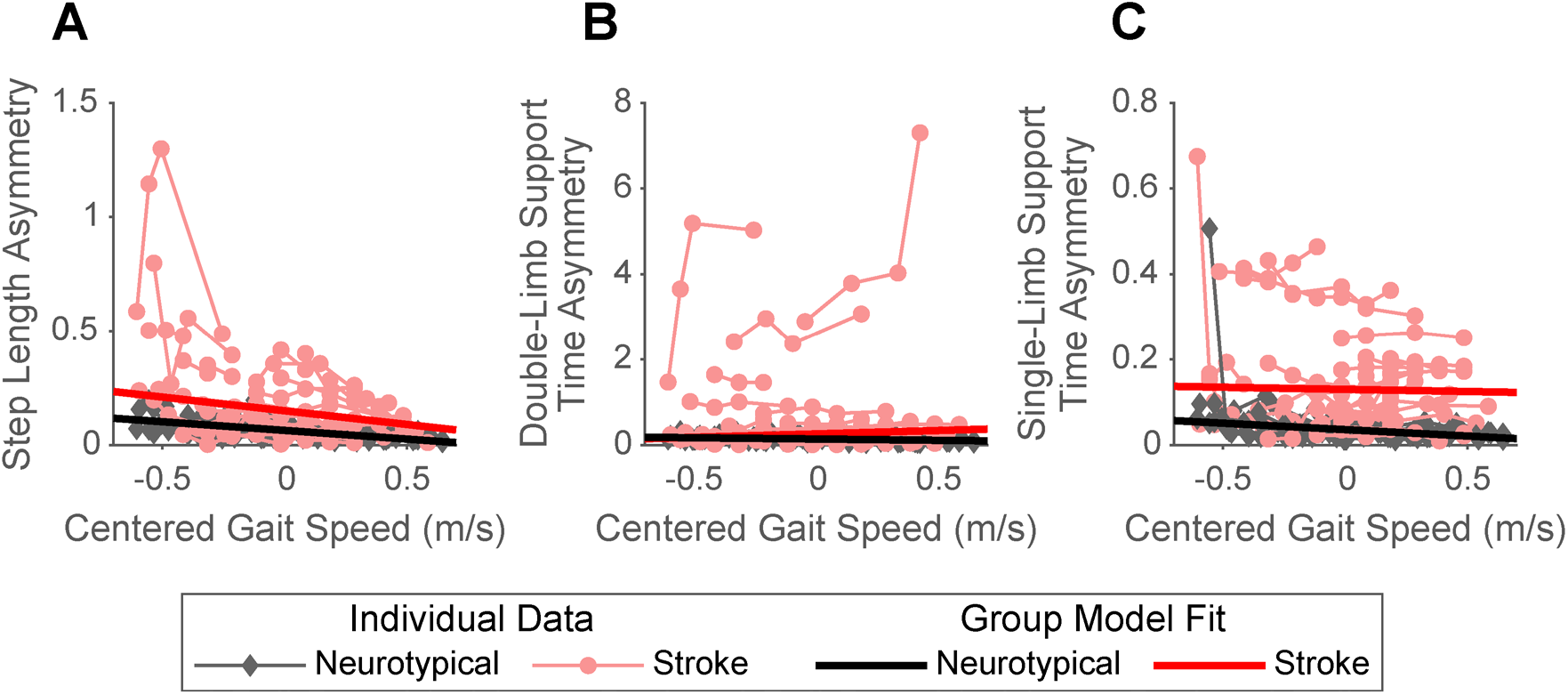
Spatiotemporal asymmetries by group across gait speeds. The figure displays the step length asymmetry **(A)**, double-limb support time asymmetry **(B)**, and single-limb support time asymmetry **(C)** of individual participants (red: post-stroke; black: neurotypical) across gait speeds. The thinner, lighter lines represent the data for an individual participant walking at 3 or 4 different gait speeds. The thicker, darker lines represent the group fits from the robust mixed-effects model.

We also found that people post-stroke exhibited more double-limb support time asymmetry relative to neurotypical adults (Figure 1B; β = 0.12, p < 0.001). Neurotypical adults decreased double-limb support time asymmetry at faster speeds (Figure 1B; β = -0.06, p < 0.001), while double-limb support time asymmetry increased in people post-stroke as demonstrated by a significant speed by group interaction (β = 0.22, p < 0.001). This means the group difference in double-limb support time asymmetry was exaggerated faster speeds.

People post-stroke also had greater single-limb support time asymmetry (Figure 1C; β = 0.09, p < 0.001) than neurotypical adults. We found a significant speed by group interaction (β = 0.02, p < 0.001), suggesting that the difference in single-limb support time asymmetry between groups increased at faster speeds. This was driven by a significant decrease in single-limb support time asymmetry exhibited by neurotypical adults with increased speeds (β = -0.03, p < 0.001).

### Kinematic Parameters

We found a significant group effect for peak swing knee flexion angle (Figure 2A; p < 0.001), with neurotypical adults having an average of 17° greater peak swing knee flexion angle than people post-stroke. Peak swing knee flexion angle of neurotypical adults increased significantly with increases in speed (β = 12.1, p < 0.001). People post-stroke also exhibited speed-dependent increases in peak swing knee flexion angle. However, the magnitude of this increase for a given change in speed was smaller than that observed in neurotypical adults (speed by group interaction: β = -8.0, p < 0.001), increasing the difference between groups at faster speeds.

**Figure 2.**
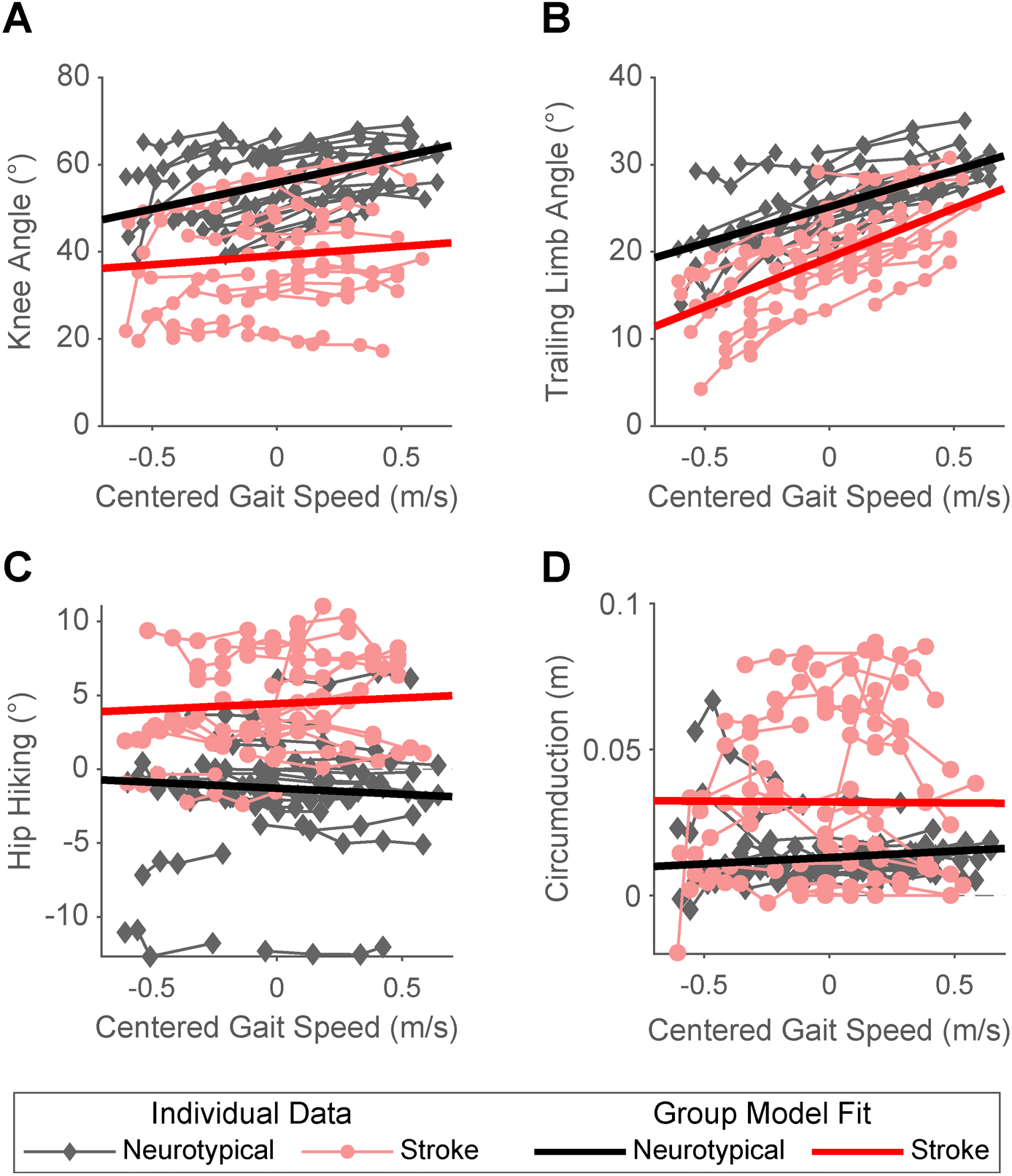
Kinematic gait parameters by group across gait speeds. The thinner, lighter lines represent the data for an individual participant (red: post-stroke; black: neurotypical) walking at 3 or 4 different gait speeds. The thicker, darker lines represent the group fits from the robust mixed-effects model. **A)** Peak swing knee angle across gait speeds. Positive values indicate knee flexion, negative values indicate knee extension. **B)** Trailing limb angle across gait speeds. **C)** Hip hiking across gait speeds. Two participants were excluded from this analysis due to missing iliac crest marker data. **D)** Circumduction across gait speeds.

People post-stroke exhibited an average of 5° less trailing limb angle compared to neurotypical adults (Figure 2B; p < 0.001). While neurotypical adults increased their trailing limb angles with faster speeds (β = 9.6, p < 0.001), people post-stroke increased their trailing limb angle more for a given increase in speed than neurotypical adults (speed by group interaction: β = 2.3, p < 0.001), decreasing the difference in trailing limb angle between the groups at faster speeds.

People post-stroke had an average of 6° greater hip hiking (Figure 2C; p < 0.001) than neurotypical adults. This difference was exaggerated at faster speeds, as people post-stroke exhibited a slight increase in hip hiking relative to neurotypical adults with increased speeds (speed: β = -0.8, p < 0.001, speed by group interaction: β = 1.6, p < 0.001). Of note, two neurotypical participants were excluded from the hip hiking analysis due to iliac crest marker occlusion.

As expected [4], we found that people post-stroke had an average of 0.02 meters greater circumduction (Figure 2D; p < 0.001) compared to neurotypical adults. There was a small increase in circumduction in neurotypical adults at faster speeds (β = 0.004, p = 0.03), but there was no significant speed by group interaction (β = -0.005, p = 0.11).

### K-means Clustering Analysis

We used k-means clustering to identify subsets of participants with similar overall gait behavior and determined whether walking at faster speeds altered the composition of these subsets. Due to missing hip hiking data, three neurotypical participants were excluded from the k-means analysis. Two participants were missing hip data at all speeds, and one participant was missing data at the fastest speed only. We chose two clusters for both the self-selected and fastest speeds using the silhouette method.

At self-selected speeds, all 47 neurotypical participants and 11/28 people post-stroke were in the first cluster (Figure 3A; diamonds). The remaining 17 people post-stroke were in the second cluster (circles). The cluster assignments were largely the same at the fastest speeds (Figure 3B), except for two participants post-stroke who switched clusters.

**Figure 3.**
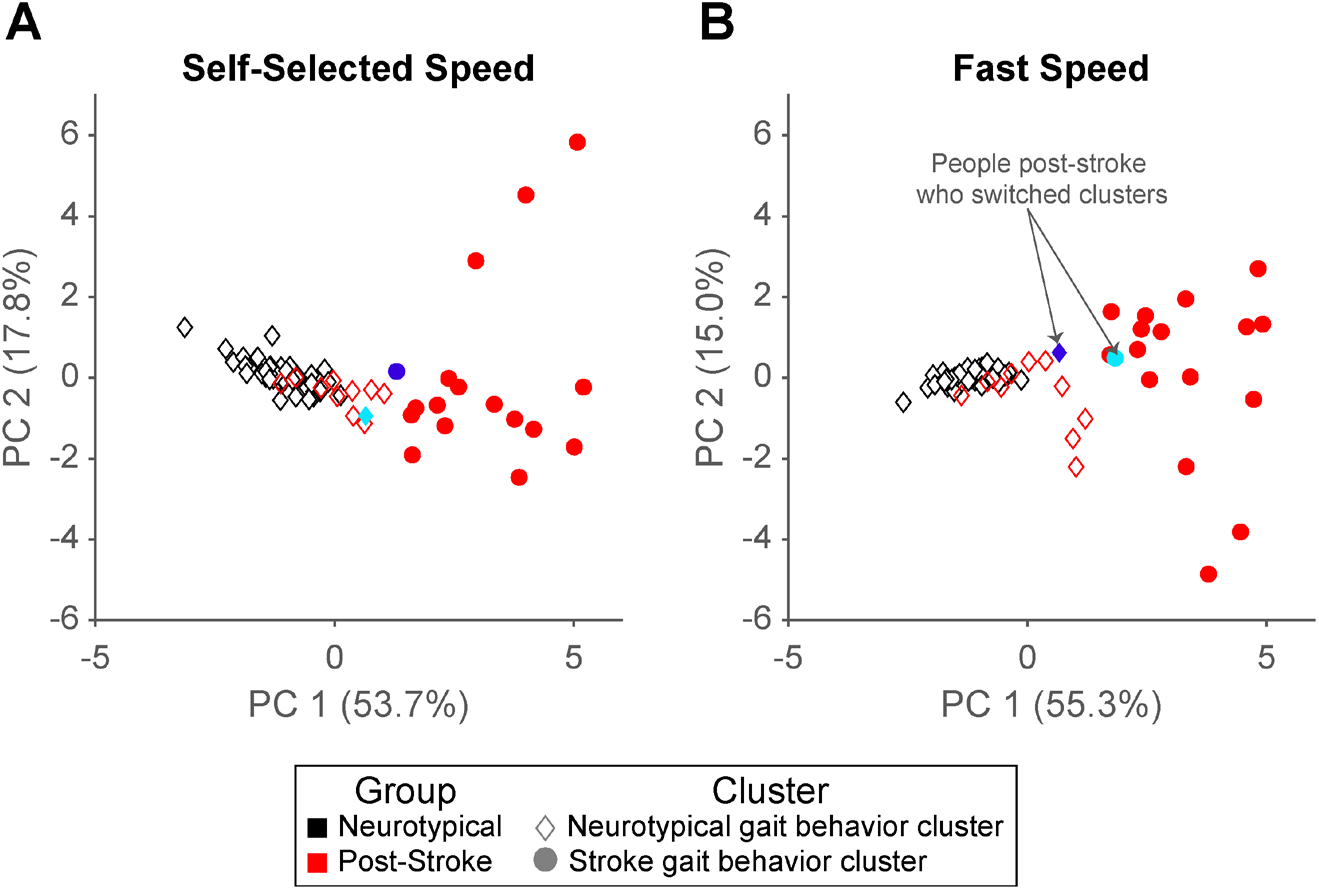
K-means clustering results across gait speeds. Individual scores for principal component 1 vs. principal component 2 are plotted to allow for visualization of two out of seven dimensions of the data included in the cluster analysis. Due to missing hip hiking data, three neurotypical participants were excluded from the k-means analysis. Marker colors represent the true group for the individual (black: neurotypical, red: post-stroke). The marker shape represents the assigned cluster (diamond: neurotypical gait behavior cluster, circle: stroke gait behavior cluster). Two participants post-stroke switched clusters at faster gait speeds. One participant moved from the neurotypical gait behavior cluster to the stroke gait behavior cluster (cyan marker), and one participant moved the opposite way (royal blue marker).

Figure 4A displays the centroids (i.e., scaled mean values) for each gait variable at both speeds for each cluster. These values indicate that the first cluster captured participants with kinematics associated with neurotypical gait behavior (e.g., larger peak swing knee flexion, smaller gait asymmetries, etc.), and the second cluster had kinematics associated with stroke gait behavior (e.g., smaller peak swing knee flexion, greater gait asymmetries, etc.). Because of this and the participant composition of the clusters, we will refer to the first cluster as the “neurotypical gait behavior cluster” and the second cluster as the “stroke gait behavior cluster.” However, it is important to note that there are individuals post-stroke in both clusters. The order of variable importance in determining the cluster assignments changed slightly from self-selected to fast speeds (Figure 4B-C; see the ranks of hip hiking and knee flexion). However, single-limb support time asymmetry was the variable most important in determining the cluster assignment regardless of speed. This is likely because participants in the stroke gait behavior cluster had much higher single-limb support time asymmetries than participants in the neurotypical gait behavior cluster at both speeds (Figure 4A; right most panel).

**Figure 4.**
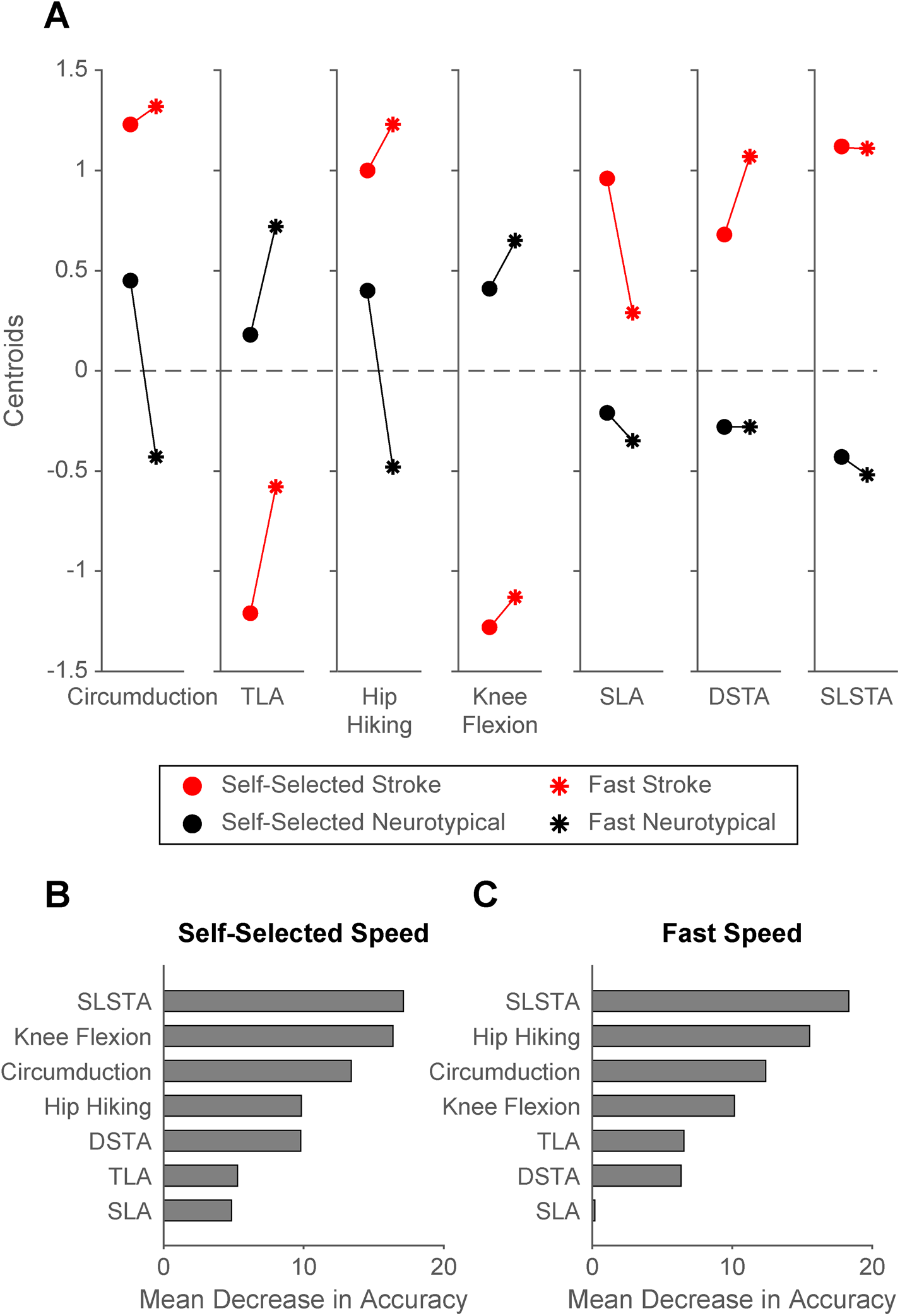
Cluster centroids **(A)**, and variable importance results **(B & C). A)** Cluster centroids (means) for each cluster at self-selected and fast speeds The red data represents the stroke gait behavior cluster, the black data represents the neurotypical gait behavior cluster. The circle markers represent the self-selected speed, and the asterisk represents the fast speed. Data were scaled (mean = 0, SD = 1) before calculating the mean value. **B)** Variable importance results at self-selected speeds. **C)** Variable importance results at fast speeds. Abbreviations: TLA, trailing limb angle; SLA, step length asymmetry; DSTA, double-limb support time asymmetry; SLSTA, single-limb support time asymmetry.

Of note, the participants post-stroke assigned to the neurotypical gait behavior cluster exhibited less lower extremity motor impairment on the Lower Extremity Fugl-Meyer than the individuals in the stroke gait behavior cluster at both speeds (Figure 5A; self-selected speed: p = 0.004; fast speed: p = 0.008). Yet, there were no differences between the participants’ post-stroke self-selected or fast speeds in each cluster (Figure 5B; self-selected speed: p = 0.59; fast speed: p = 0.43).

**Figure 5:**
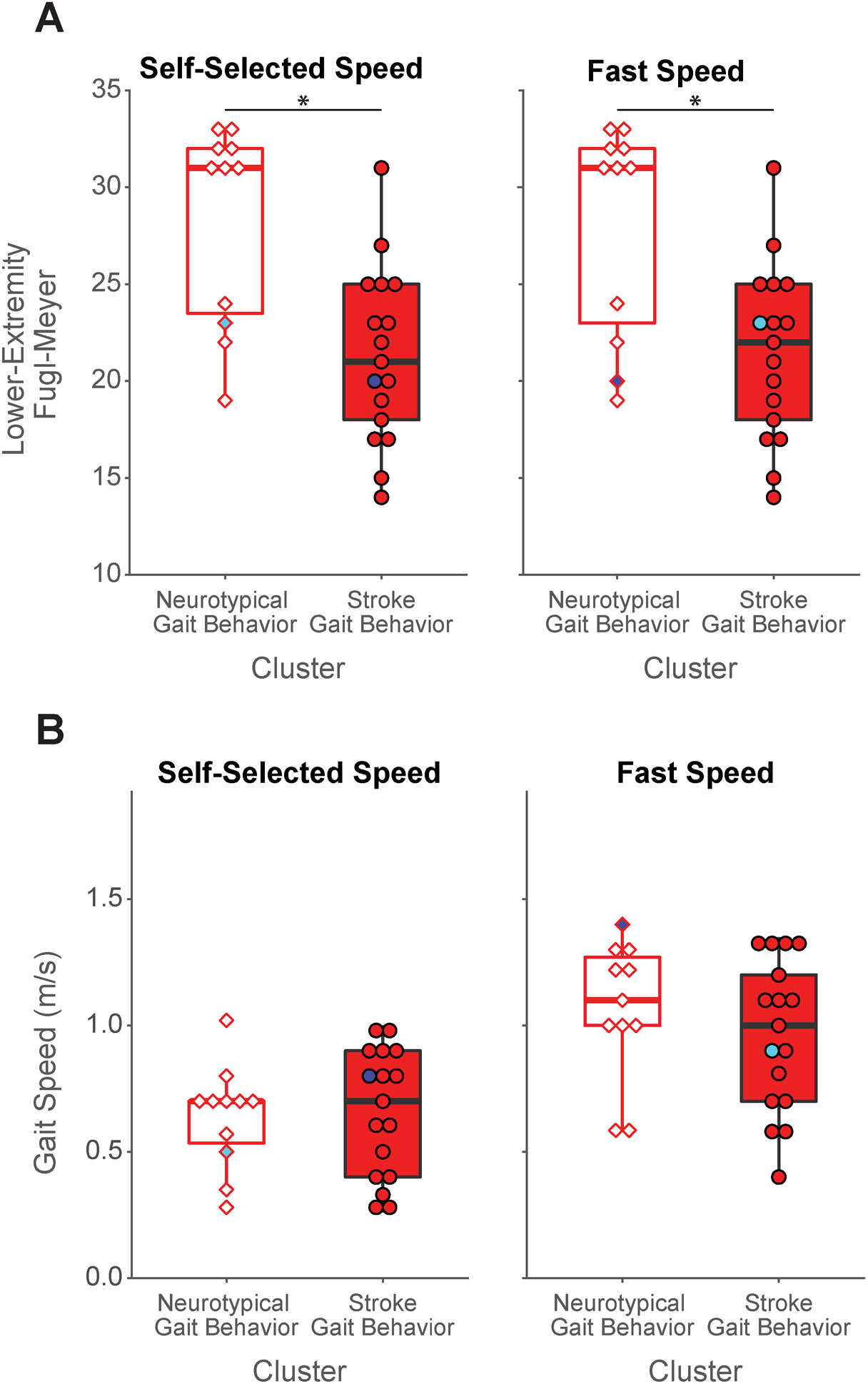
Lower Extremity Fugl-Meyer scores **(A)** and gait speeds **(B)** of participants post-stroke in each cluster. The white-filled boxplots represent participants post-stroke in the neurotypical cluster. The red-filled boxplots represent participants post-stroke in the stroke gait behavior cluster. Two participants switched clusters at faster gait speeds. One participant moved from the neurotypical gait behavior cluster to the stroke gait behavior cluster (cyan marker), and one participant moved the opposite way (royal blue marker).

Two participants post-stroke were classified into a different cluster when walking at faster speeds. One moved from the stroke gait behavior cluster at self-selected speed to the neurotypical gait behavior cluster at the fast speed (Figures 3A and 3B; royal blue symbol). The other switched from the neurotypical gait behavior cluster at self-selected speed to the stroke gait behavior cluster at the fast speed (Figures 3A and 3B; cyan symbol).

Finally, we found that the kinematics of the individuals in the stroke gait behavior cluster were more variable and different than those in the neurotypical gait behavior cluster at the fast speed relative to self-selected speed. The within sum of squares (capturing within-cluster variability) of the neurotypical gait behavior cluster decreased at the fast speed (self-selected speed: 107.3; fast speed: 92.2) while it increased in the stroke gait behavior cluster (self-selected speed: 156.5; fast speed: 167.7). The two clusters also moved further apart at fast speeds, demonstrated by the increase in between sum of squares (self-selected speed: 187.7; fast speed: 207.6). It is important to note that the variability within and between clusters is not well-represented in Figure 3 because the data is plotted in the principal components space, with only two out of seven dimensions plotted.

## DISCUSSION

This study aimed to determine the effect of fast walking on gait kinematics in people post-stroke relative to neurotypical adults. Consistent with our hypothesis, speed-dependent changes in peak swing knee flexion, hip hiking, double-limb support time asymmetry, and single-limb support time asymmetry lead to a larger between-group difference at faster speeds. Contrary to our hypothesis, faster walking did not affect between-group differences in circumduction and reduced between-group differences in step length asymmetry and trailing limb angle. We found two distinct clusters when we included all kinematic metrics in clustering analyses during self-selected and fast walking to understand the effect on overall gait behavior. One cluster represented neurotypical gait behavior and was composed of all the neurotypical participants (n = 47) and a sub-group of the participants post-stroke (n = 11/28). The other characterized post-stroke gait behavior and contained only participants post-stroke (n = 17/28). At fast speeds, cluster assignments largely did not change, but the clusters were further differentiated. When all metrics were considered simultaneously, at fast speeds, the overall gait pattern of a subgroup of people post-stroke became more distinct from participants (post-stroke and neurotypical) who demonstrated an overall gait pattern consistent with neurotypical adults. These findings indicate that the biomechanical benefit (i.e., the change of metric in the same direction as that observed in neurotypical adults) of fast walking post-stroke varies across individual gait metrics and in a sub-group of participants post-stroke (n = 17/28) these benefits did not lead to an overall gait pattern more similar to neurotypical adults. This suggests that there is an opportunity to potentiate speed-dependent biomechanical changes by coupling fast walking with other interventions.

Our evaluation of individual gait metrics found that the relative speed-dependent changes in four of the seven aligned with our hypothesis that between-group differences would be larger at faster speeds. For example, walking at faster speeds increased the absolute magnitude of peak swing knee flexion in the participants post-stroke. Yet, the magnitude of speed-dependent increase was much smaller than observed in neurotypical adults. This small speed-dependent change in swing knee flexion, relative to that observed in neurotypical adults, likely explains the persistence of compensatory gait deviations, such as hip hiking and circumduction, at faster speeds [27,28].

A few mechanisms may explain this smaller speed-dependent change in knee flexion in our participants post-stroke. First, people post-stroke are well-known to exhibit reduced neuromuscular complexity (i.e., abnormal muscle synergies, merged motor modules) in the lower extremity during walking [17–19] that may limit their capacity for large changes in knee flexion during swing phase. In addition to this, studies of the upper extremity post-stroke demonstrate that raising the demands of a movement task can increase the expression of abnormal synergistic movements [29,30]. This suggests that fast walking may increase the expression of lower extremity synergies that may limit knee flexion, similar to that observed during maximal hip extension on a dynamometer [20], particularly if participants are attempting to increase their trailing limb angle to increase propulsion to walk at faster speeds (see trailing limb angle discussion below) [31]. However, a previous study of neuromuscular complexity in participants post-stroke during walking demonstrated that the organization of motor modules does not change during fast walking [32]. This is indirectly supported by our finding that people post-stroke assigned to the neurotypical gait behavior cluster at self-selected speeds were not assigned to the stroke gait behavior cluster at fast speeds.

Another potential explanation for the smaller speed-dependent changes in peak swing knee flexion observed in people post-stroke is increased spastic activity of the quadriceps on the paretic limb [33–35]. There is a strong relationship between the magnitude of peak swing knee flexion and measures of stretch reflex hyperactivity in the quadriceps muscles in people post-stroke [35,36]. Moreover, an imposed stretch to the hip flexors post-stroke leads to a hyper-activation of the stretch reflex in the quadriceps that increases with faster stretch velocities [35]. Walking at faster speeds on a treadmill with increasing trailing limb angles will naturally lead to a faster stretch of the hip flexors during late stance than walking slower. This, in combination with potential abduction-dependent reflex activation [37], may cause involuntary hyper-activation of the quadriceps, contributing to smaller speed-dependent changes in peak swing knee flexion. Lastly, reduced peak swing knee flexion post-stroke has also been associated with diminished paretic limb propulsion [38,39].

In contrast to the speed-dependent exaggeration of between-group differences described above, speed-dependent reductions in step length asymmetry within this single session led to smaller between-group differences at faster speeds. This suggests that the biomechanical byproduct of reduced step length asymmetry during fast walking post-stroke may be sufficient to reduce the difference relative to neurotypical gait. Yet, people post-stroke have consistently demonstrated no significant change in step length asymmetry following long-term fast walking interventions [14,40].

People post-stroke also exhibited larger increases in trailing limb angle for a given change in speed relative to neurotypical adults, decreasing the magnitude of the difference between the groups at faster speeds. This finding is likely related to the increased propulsion needed to walk faster. Propulsion during walking can be altered by increasing plantar-flexor moment and trailing limb angle [41]. However, people post-stroke tend to increase propulsion during fast walking through increases in trailing limb angle, without changing plantar-flexor moment [31]. This suggests that people post-stroke were modulating propulsion primarily through changes in trailing limb angle, while neurotypical participants were using both strategies to meet the increased propulsion demands of fast walking.

Given that the relative speed-dependent effects varied across individual gait metrics, we used k-means clustering to consider all gait metrics simultaneously and gain insight into the relative effect of speed on overall gait behavior post-stroke. This revealed two clusters at self-selected and fast speeds. One cluster characterized gait behavior typically found in neurotypical adults, and the other characterized gait behavior typically found in people post-stroke. Our analysis of variable importance in participants’ cluster assignment found that magnitudes of single-limb stance time asymmetry, knee flexion, and circumduction were the most influential in determining the clusters at self-selected speed (Figure 4C). This changed slightly at the fast speeds – with single limb stance time asymmetry, hip hiking, and circumduction being the top three determinants of cluster classification (Figure 4D). Approximately 40% of the participants post-stroke in our sample (n = 11/28) were classified as part of the neurotypical gait behavior cluster at both speeds. This indicates that this subset of post-stroke participants walked with an overall gait pattern more similar to neurotypical than the other participants post-stroke. The centroids (i.e., scaled mean values) for each gait variable for each cluster (Figure 4A) and the order of variable importance analysis (Figures 4B and 4C) suggest that these participants exhibited less single-limb stance time asymmetry, hip hiking, and circumduction and more knee flexion than the post-stroke participants assigned to the other cluster. This subset of participants also had significantly less lower extremity motor impairment (Figure 5A) than those classified in the stroke gait behavior cluster. This highlights an opportunity for future work to 1) determine other clinical characteristics that may contribute to heterogeneity in overall gait behavior post-stroke and 2) identify subgroups of people post-stroke that may benefit more from combining fast walking with an approach that strategically targets gait biomechanics.

Fast walking alone did not lead to biomechanical changes that were large or consistent enough to cause participants post-stroke to change cluster assignments. Only two participants post-stroke were assigned to the opposite cluster when walking at faster speeds – one to the neurotypical cluster and one to the post-stroke cluster (royal blue and cyan points, respectively in Figures 3A and 3B). Fast walking also did not cause the clusters to move closer together, as we might expect if overall gait behavior more similar to neurotypical were a byproduct of fast walking. Instead, we found that when walking at fast speeds, participants in the stroke gait behavior cluster exhibited an overall gait pattern that was more different and more variable than those of the participants (both post-stroke and neurotypical) in the neurotypical gait behavior cluster.

### Limitations

This study has a few limitations. First, the data were collected at three sites with different motion capture set-ups and study protocols. Because the participants collected at each site were not equally balanced between groups (i.e., one site collected data from only stroke participants, one site collected data from only neurotypical adults, and one site collected data from both), we could not account for this in our model. Next, we did not have access to ground reaction force or EMG data. Therefore, speed-dependent differences between groups in kinetic and muscular activity outcomes are unclear. Finally, we only had data from the paretic limb for 20/28 of the participants post-stroke. Consequently, we could not study the effect of fast walking on non-paretic limb kinematics.

## CONCLUSIONS

This study demonstrated that the biomechanical changes resulting from fast walking post-stroke, relative to neurotypical adults, vary across individual gait metrics. We found two distinct clusters representative of neurotypical gait behavior and stroke gait behavior, which became more distinct at fast speeds. What remains to be seen is if it is possible to combine fast walking with an approach, such as gait biofeedback, to strategically target gait metrics that have smaller speed-dependent changes. This would allow for a single intervention to address both activity limitations and kinematic impairments in post-stroke gait rehabilitation, which could have additive or synergistic effects on the rehabilitation of walking dysfunction post-stroke.

## ACKNOWLEDGEMENTS

We thank Dr. Natalia Sánchez for her thoughtful comments and suggestions on the k-means clustering analysis and Dr. Russell Johnson for his careful review and comments on the manuscript.

