## Supplemental Table 1 for "Speed-dependent biomechanical changes vary across individual gait metrics post-stroke relative to neurotypical adults"

**Supplemental Table 1:** Demographics of neurotypical adults.

| Data Set | Age (years) | Sex | Speed 1 (m/s) | Speed 2 (m/s) | Speed 3 (m/s) | Speed 4 (m/s) |
|---|---|---|---|---|---|---|
| Finley and Bastian | 68 | M | 0.77 | 0.96 | 1.15 | 1.24 |
| Finley and Bastian | 63 | F | 0.46 | 0.57 | 0.68 | 0.81 |
| Finley and Bastian | 57 | M | 0.21 | 0.26 | 0.31 | 0.56 |
| Finley and Bastian | 56 | F | 0.48 | 0.61 | 0.72 | 1.00 |
| Finley and Bastian | 43 | M | 0.82 | 1.02 | 1.22 | 1.35 |
| Finley and Bastian | 52 | M | 0.26 | 0.33 | 0.40 | - |
| Finley and Bastian | 55 | M | 0.28 | 0.35 | 0.42 | 0.60 |
| Finley and Bastian | 65 | M | 0.22 | 0.28 | 0.34 | 0.57 |
| Fukuchi et al. | 59 | M | 0.64 | 0.82 | 0.99 | 1.17 |
| Fukuchi et al. | 57 | F | 0.75 | 0.95 | 1.16 | 1.36 |
| Fukuchi et al. | 55 | M | 0.78 | 0.99 | 1.2 | 1.41 |
| Fukuchi et al. | 50 | M | 0.58 | 0.80 | 1.02 | 1.23 |
| Fukuchi et al. | 71 | M | 0.57 | 0.78 | 1.00 | 1.21 |
| Fukuchi et al. | 58 | F | 0.92 | 1.08 | 1.24 | 1.4 |
| Fukuchi et al. | 63 | F | 0.61 | 0.78 | 0.95 | 1.11 |
| Fukuchi et al. | 61 | F | 0.75 | 0.95 | 1.15 | 1.36 |
| Fukuchi et al. | 63 | M | 0.89 | 1.08 | 1.27 | 1.46 |
| Fukuchi et al. | 62 | M | 0.42 | 0.48 | 0.73 | 0.89 |
| Fukuchi et al. | 68 | M | 0.52 | 0.71 | 0.90 | 1.10 |
| Fukuchi et al. | 63 | M | 0.80 | 1.02 | 1.24 | 1.46 |
| Fukuchi et al. | 73 | M | 0.61 | 0.78 | 0.95 | 1.11 |
| Fukuchi et al. | 56 | F | 0.80 | 1.02 | 1.24 | 1.46 |
| Fukuchi et al. | 84 | M | 0.40 | 0.54 | 0.69 | 0.84 |
| Fukuchi et al. | 68 | F | 0.74 | 0.94 | 1.14 | 1.34 |
| Fukuchi et al. | 55 | F | 0.47 | 0.64 | 0.82 | 0.99 |
| Fukuchi et al. | 63 | F | 0.36 | 0.49 | 0.63 | 0.76 |
| Fukuchi et al. | 25 | M | 0.49 | - | - | 1.03 |
| Fukuchi et al. | 22 | F | 0.5 | - | - | 0.69 |
| Fukuchi et al. | 33 | M | 0.39 | - | - | 1.12 |
| Fukuchi et al. | 24 | M | 0.71 | - | - | 0.91 |
| Fukuchi et al. | 28 | M | 0.9 | - | - | 1.1 |
| Fukuchi et al. | 25 | M | 0.9 | - | - | 1.28 |
| Fukuchi et al. | 24 | F | 0.61 | - | - | 1.1 |
| Fukuchi et al. | 36 | M | 0.76 | - | - | 0.97 |
| Fukuchi et al. | 25 | F | 0.82 | - | - | 0.99 |
| Fukuchi et al. | 31 | F | 0.76 | - | - | 1.37 |
| Fukuchi et al. | 32 | M | 0.92 | - | - | 1.31 |
| Fukuchi et al. | 24 | F | 0.58 | - | - | 0.9 |
| Fukuchi et al. | 30 | M | 1.02 | - | - | 1.33 |
| Fukuchi et al. | 31 | F | 0.78 | - | - | 0.62 |
| Fukuchi et al. | 23 | M | 0.85 | - | - | 1.31 |
| Fukuchi et al. | 31 | M | 1.1 | - | - | 1.27 |
| Fukuchi et al. | 28 | M | 0.55 | - | - | 1.16 |
| Fukuchi et al. | 28 | F | 0.5 | - | - | 0.88 |
| Fukuchi et al. | 29 | M | 1.03 | - | - | 1.21 |
| Fukuchi et al. | 21 | F | 0.9 | - | - | 1.09 |
| Fukuchi et al. | 22 | M | 0.99 | - | - | 1.21 |
| Fukuchi et al. | 25 | F | 0.73 | - | - | 1.12 |
| Fukuchi et al. | 28 | F | 0.84 | - | - | 1.06 |
| Fukuchi et al. | 37 | M | 1.02 | - | - | 1.17 |

Neurotypical adults from Fukuchi et al (2018) were speed-matched to the participants post-stroke in the Tyrell et al (2011) data set. The shaded region represents the younger, neurotypical adults used only in the k-means clustering analysis.
